# Reciprocal regulation of the H_3_ histamine receptor in Rett syndrome and *MECP2* Duplication syndrome: implications for therapeutic development

**DOI:** 10.1101/2025.10.30.685636

**Authors:** Kelly Weiss, Sheryl A. D. Vermudez, Geanne Freitas, Shalini Dogra, Mac J. Meadows, Rocco G. Gogliotti, Colleen M. Niswender

**Affiliations:** Department of Pharmacology and Warren Center for Neuroscience Drug Discovery, Vanderbilt University, Nashville, TN 37232 (KW, SADV, GF, SD, MJM, RGG, CMN); Vanderbilt Institute of Chemical Biology, Vanderbilt University, Nashville, TN 37232 (CMN); Vanderbilt Brain Institute, Vanderbilt University, Nashville, TN 37232 (CMN); Vanderbilt Kennedy Center, Vanderbilt University Medical Center, Nashville, TN 37232 (CMN)

## Abstract

Rett syndrome (RTT) and *MECP2* Duplication syndrome (MDS) are disorders caused by reciprocal decreases and increases in the expression of the transcriptional regulator, *Methyl CpG Binding Protein 2* (*MeCP2*). We previously performed an mRNA expression profiling study of the temporal cortex region from patients diagnosed with RTT and corresponding age, postmortem interval, and sex-matched controls. These studies identified a significant reduction in the expression of the histamine H_3_ receptor (*HRH3*). In the current manuscript, we expanded this H_3_ receptor profiling to additional RTT patient brain samples representing distinct *MECP2* mutations and confirmed significantly reduced levels of H_3_ receptor expression in the majority of patients compared to controls. Using mouse models of RTT and MDS, we observed antiparallel changes in H_3_ receptor expression across various brain areas, with *Hrh3* expression being reduced in RTT model animals and increased in a mouse model of MDS. We then evaluated both a small molecule agonist of the H_3_ receptor, (*R*)-α-methylhistamine (RAMH), and the H_3_ receptor inverse agonist, pitolisant (Wakix®), in RTT and MDS models, respectively, to determine impacts on phenotypes in these disease models. Our results show that RAMH significantly impacted an anxiety phenotype in mice modeling RTT (*Mecp^Null/+^*), but pitolisant had no effect on the behaviors examined here in MDS animals (*MECP2^Tg1^*).

## Introduction

Rett syndrome (RTT) is caused by mutations in the X-linked gene encoding Methyl CpG Binding Protein 2 (MeCP2), a transcriptional regulator of both local and global gene expression patterns (Amir, Van den Veyver et al. 1999, Chahrour, Jung et al. 2008, Ben-Shachar, Chahrour et al. 2009). Although RTT can occur in either sex, RTT patients are predominantly female and develop normally during their first 6 to 18 months of life; they then undergo rapid developmental regression, resulting in impairments in language and motor skills, breathing abnormalities such as apneas, and the emergence of seizures (Neul, Kaufmann et al. 2010). In addition to RTT, patients can also exhibit duplication or triplication of the *MECP2* gene, resulting in *MECP2* duplication syndrome (MDS), a disorder characterized by numerous abnormal symptoms such as hypotonia, seizures, intellectual disability, and recurrent infections (Ramocki, Peters et al. 2009, Ramocki, Tavyev et al. 2010, Collins and Neul 2022). One symptomatic treatment currently available for RTT is the recently approved compound trofinetide (Daybue®) (Neul, Percy et al. 2023); other strategies are aimed specifically at MeCP2 and include gene therapy, DNA and RNA editing, X-chromosome reactivation, and pharmacological read-through approaches (reviewed in (Collins and Neul 2022, Palmieri, Pozzer et al. 2023); NCT05898620, NCT06152237). There are currently no approved therapeutic options specifically for the treatment of MDS.

The availability of expression profiling data from the brains of RTT patients provides an opportunity to identify or confirm therapeutic targets or clarify biological mechanisms important for RTT that begin from a translational perspective. We previously performed an RNA expression profiling study of the temporal cortex region using autopsy samples from patients diagnosed with RTT and corresponding age, postmortem interval, and sex-matched controls (Gogliotti, Fisher et al. 2018). These studies identified a number of G protein-coupled receptors (GPCRs) that were changed in expression in RTT patients, including several metabotropic glutamate (mGlu) and muscarinic receptor candidates (Gogliotti, Fisher et al. 2018). Further evaluation of these RNA sequencing data revealed significantly reduced expression of the histamine H_3_ receptor (*HRH3*) in brain samples from the cerebellum of RTT patients. As shown here, expansion of this profiling set into a larger cohort of temporal cortical samples from 14 control and 29 RTT patients revealed significant decreases in the expression of *HRH3* mRNA in the majority of RTT patients compared to controls. Additionally, we verified decreased *Hrh3* mRNA expression in several brain areas of *Mecp2^Null/+^* mice, a commonly used model of RTT (Guy, Hendrich et al. 2001). In parallel, we also confirmed increased levels of *Hrh3* mRNA in brains from mice modeling MDS (*MECP2^Tg1^*, (Collins, Levenson et al. 2004)). These observations correlate with findings from other studies showing decreases in mouse *Hrh3* expression in male mice lacking Mecp2 (*Mecp2^Null/y^*), as well as reciprocal increases in expression in *Mecp2Tg1* animals (Chahrour, Jung et al. 2008, Ben-Shachar, Chahrour et al. 2009, Renthal, Boxer et al. 2018). Additionally, a study profiling the occipital cortex from four female RTT patients demonstrated a significant decrease in the expression of *HRH3* mRNA in cells expressing an MeCP2 mutant (Supplemental Table 7 in (Renthal, Boxer et al. 2018)). Recently, a profiling study of RTT patients also reported decreased levels of histidine, the precursor for histamine, in the plasma of RTT patients compared to unaffected siblings (Neul, Skinner et al. 2020).

Histaminergic projections originate in the tuberomammillary nucleus of the hypothalamus and H_3_ receptors are found throughout the brain with particular abundance in the cortex, thalamus, and nucleus accumbens (Leurs, Bakker et al. 2005, Nieto-Alamilla, Marquez-Gomez et al. 2016). Activation of the presynaptic H_3_ receptor modulates not only the release of histamine, but also other neurotransmitters such as dopamine, serotonin, GABA, and acetylcholine (Brown, Stevens et al. 2001, Galici, Boggs et al. 2009). H_3_ receptors expressed on synapses projecting to the nucleus accumbens (NAc) regulate anxiety phenotypes and repetitive behaviors by, among other mechanisms, decreasing presynaptic release of glutamate (Zhang, Peng et al. 2020). Additionally, mice fed a low-histidine diet for two weeks exhibit lower levels of brain histamine and release less histamine in the hypothalamus after stimulation; these animals also display increases in anxiety-related behaviors in elevated maze and light-dark transition tasks (Yoshikawa et al., 2014).

Based on our expression profiling, we evaluated the activity of a small molecule agonist of the H_3_ receptor, (*R*)-α-methylhistamine (RAMH), on several behaviors in *Mecp2^Null/+^* mice; additionally, we tested the hypothesis that the clinically available H_3_ receptor antagonist/inverse agonist pitolisant (Wakix®) might be effective in correcting phenotypes in MDS animals. Our results show that RAMH enhanced an anxiolytic effect in *Mecp2^Null/+^* animals; however, pitolisant was without effect in *MECP2^Tg1^* animals in the behavioral assays profiled here. Overall, these results suggest that there may be therapeutic utility in increasing H_3_ receptor activity to treat specific domains of RTT.

## Materials and Methods

### Human Brain Sample Profiling

Human samples were obtained from the National Institutes of Health NeuroBioBank (neurobank.nih.gov) under Public Health Service contract HHSN-271-2013-00030. The tissues were post-mortem and fully de-identified, and as such are classified as exempt from human subject research regulations. The demographics of autopsy samples used in these studies are shown in Table 1. In this study, we only used samples with verified mutations in *MECP2* and did not include samples that were negative for *MECP2* mutation or for which the *MECP2* genotype could not be conclusively defined.

**Table 1.**
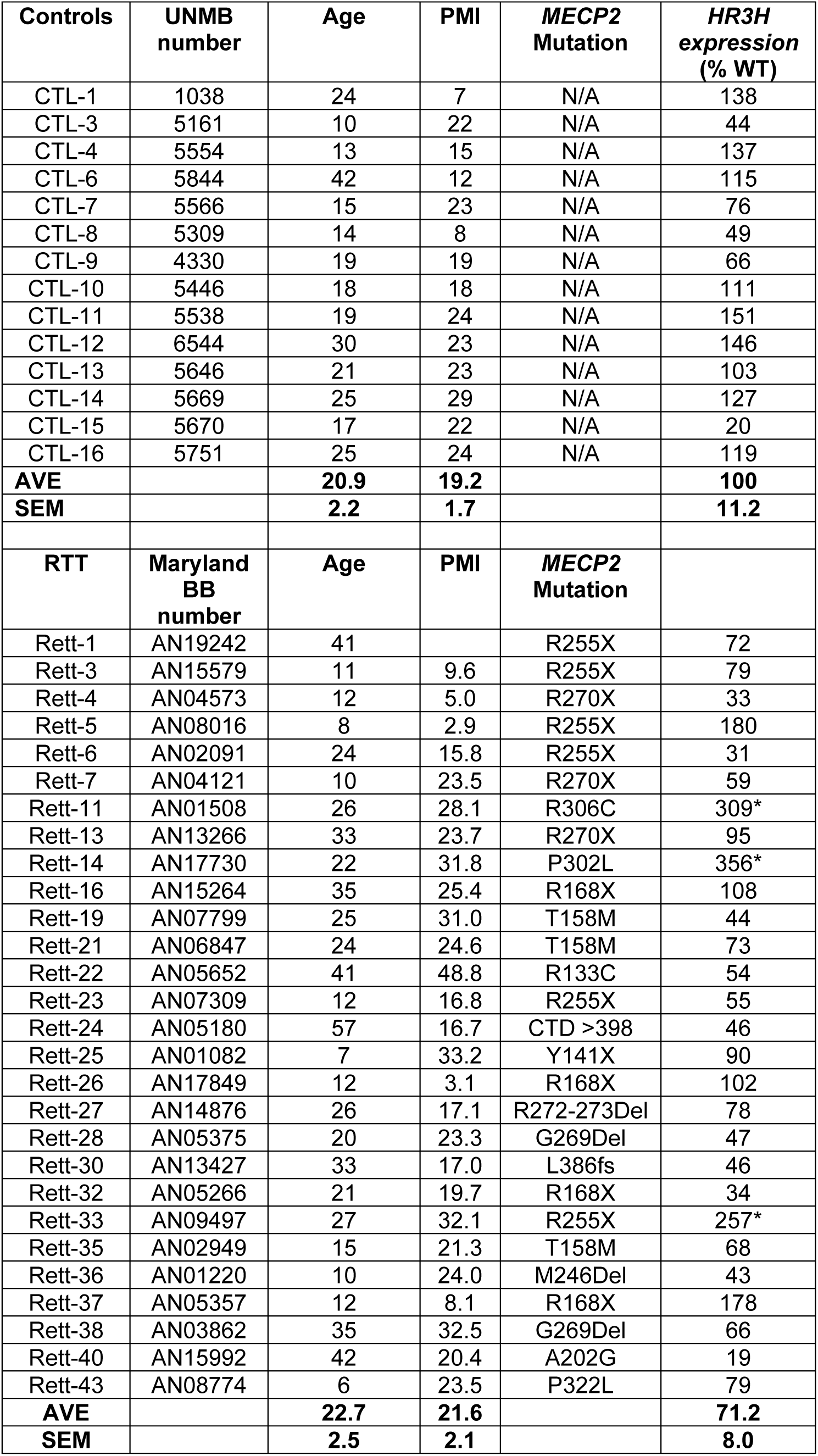
Demographics of human patient samples. Temporal cortex samples listed below were used in Figure 1C. Samples determined to be outliers by ROUT analysis are shown with a * designation.

### RNA-cDNA Preparation

Approximately 1 g of human tissue was impact dissociated under dry ice and then pulverized using mortar and pestle under liquid nitrogen. Total RNA was prepared from 100 to 200 mg of tissue using standard Trizol-chloroform methodology. Total RNA was purified and DNase-treated using an RNeasy kit (Qiagen, Hilden, Germany). RNA quality was confirmed using BioCube and BioAnalyzer instrumentation by the Vanderbilt Technologies for Advanced Genomics core (Vanderbilt University Medical Center). cDNA was synthesized from 2 μg total RNA with the SuperScript VILO kit (Thermo Fisher, Waltham, MA). Samples from the cortex, striatum, cerebellum, and hippocampus were prepared from mice using identical methodology.

### Quantitative RT-PCR (qRT-PCR)

qRT-PCR (CFX96, Bio-Rad, Vanderbilt University Medical Center MCBR core) using 50 ng/9 μL cDNA from the cortex, hippocampus, striatum and cerebellum samples from *Mecp2^Null/+^* and *MECP2^Tg1^* mice was run in duplicate using PowerUp™ SYBR™ Green Master Mix (ThermoFisher cat no. A25742) with the following primers (5’ to 3’): human H_3_ receptor (forward: TCTTCCTGCTCAACCTCGCCAT, reverse: ACTACCAGCCACAGCTTGCAGA, 127 bp), mouse H_3_ receptor (forward: CTTCCTCGTGGGTGCCTTC, reverse: CAGCTCGAGTGACTGACAGG, 179 bp), and human/mouse *GAPDH*/*Gapdh* (forward: CGACTTCAACAGCAACTCCC, reverse: GCCGTATTCATTGTCATACCAGG, 106 bp). All primer sequences were designed using Primer3 (SYBR Green) and constructed by Sigma through the Vanderbilt University Medical Center MCBR core. Ct values for each sample were normalized to *GAPDH*/*Gapdh* expression and analyzed using the delta−delta Ct method (Livak and Schmittgen 2001). Each value was compared to the average delta-Ct value acquired for wild-type controls and calculated as percent-relative to the average control delta-Ct.

### Animals

All animals used in the present study were group housed with food and water given ad libitum and maintained on a 12hr light/dark cycle. Animals were cared for in accordance with the National Institutes of Health Guide for the Care and Use of Laboratory Animals. All studies were approved by the Vanderbilt Institutional Animal Care and Use Committee and took place during the light phase. *Mecp2^Null/+^* (B6.129P2(C)-*Mecp2^tm1.1Bird^*/J, stock no. 003890) mice were obtained from The Jackson Laboratory (Bar Harbor, ME, USA) and maintained on a C57BL/6J background by breeding *Mecp2^Null/+^* animals with WT C57BL/6J mice (The Jackson Laboratory, stock no. 000664). As a reflection of the predominantly female RTT patient population, female *Mecp2^Null/+^* mice were utilized and aged to at least 20 weeks of age prior to experiments. *MECP2^Tg1/o^* mice (C57BL/6J background) were generously shared by Dr. Jeffrey Neul (Vanderbilt University Medical Center) and were bred with C57BL/6J animals. Both sexes of mice were used between 6-10 weeks of age.

### Drugs

RAMH, the *R*-enantiomer of α-methylhistamine dihydrobromide (H_3_ receptor agonist) was purchased from Tocris Bioscience (catalog number 0569, Bristol, UK) and pitolisant hydrochloride was purchased from AdooQ BioScience (catalog number 15213, Irvine, CA). All drugs for behavioral experiments were diluted in 0.9% saline.

### Behavioral Assays

All behavioral experiments were conducted at predicted symptomatic ages (20-25 week-old female *Mecp2^Null/+^* mice and 8-12 weeks for male and female *MECP2^Tg1^* animals, as well as corresponding littermate controls) at the Vanderbilt Mouse Neurobehavioral Lab Core. All experiments were preceded by intraperitoneal (i.p.) injections: vehicle (0.9% sterile saline), RAMH (45 mg/kg, 15 min pretreatment) or pitosilant hydrochloride (10 mg/kg, 30 min pretreatment). Quantification was performed either by a researcher blinded to the genotype and/or by automated software.

### Open Field

Mice were placed in the activity chamber for 60 min. Exploratory and locomotor activity was quantified using Activity software by assessing beam breaks in the X, Y and Z axis (Med Associates Inc).

### Elevated Zero Maze

The elevated zero maze has two regions that are closed (contain walls) and two regions that are open (no walls). Mice were allowed to freely explore the maze for five minutes under full light conditions (∼ 700 lux in the open regions, ∼ 400 lux in the closed regions). The percentage of time spent in the open versus closed regions and number of region-transitions was measured. The assay was tracked by a mounted camera and analyzed by automated AnyMaze software (Stoelting, Wood Dale, IL, USA).

### Hindlimb clasping

*Mecp2^Null/+^* female mice show a clasping phenotype that becomes apparent in our colony at 16 weeks of age (Gogliotti, Senter et al. 2017). In mice, hind limb clasping was assessed by suspending a mouse by its tail and video recording clasping dynamics for a period of 1 minute, which were scored by a blinded reviewer.

### Fear Conditioning

On both the training (Day 1) and test days (Day 2), mice were habituated to the testing room for 1 hour prior to the start of the behavioral assay. On Day 1, mice received either vehicle, RAMH or pitolisant. Following treatment, mice were placed into the operant chamber equipped with a shock grid (Med Associates Inc.) and a 10% vanilla odor cue. After a 3-minute acclimation period, mice received 2x foot shocks (0.7 mA, 2 seconds duration) separated by a 60 second interval, followed by an additional 30 second recovery period in the chamber. On Day 2, 24 hours later, mice were returned to the same chamber for 3 minutes in the presence of 10% vanilla odor cue; however, no foot shocks were administered. Freezing behavior was recorded and quantified using Video Freeze software (Med Associates Inc., St. Albans, VT, USA).

### Statistical Analyses

Statistics were carried out using GraphPad Prism and Excel (Microsoft). All data shown represent mean ± SEM. Statistical significance between genotypes was determined using Student’s t-test with or without Welch’s correction or 1- or 2-way ANOVA with Tukey’s post-hoc tests. Statistical tests and results of statistical analyses are specified in each figure legend.

## Results

### Histamine receptor H_3_ expression is reduced in brain samples from RTT patients and *Mecp2^Null/+^* mice

We previously performed an RNA sequencing study using nine cortical and six cerebellar brain samples from patients with either R168X, R255X, or R270X mutations in *MECP2,* as well as control samples matched for age, brain region, postmortem interval, and sex (Gogliotti, Fisher et al. 2018). These studies identified a significant reduction in mRNA expression of the histamine H_3_ receptor (*HRH3*) in the cerebellum of RTT patients versus controls (control abundance, 8.040, RTT abundance, 4.789; fold change -0.747, *p=0.011, (Gogliotti, Fisher et al. 2018)). We confirmed that *HRH3* mRNA expression was decreased in these cerebellar samples using qRT-PCR (**Figure 1A**); although not significant in the initial RNA sequencing study, we also observed a significant decrease in *HRH3* expression in the motor cortex using qRT-PCR (**Figure 1B).** We then expanded these studies to samples from the temporal cortex of a larger cohort of RTT patients representing different *MECP2* mutations (**Table 1**). We observed a range of expression levels; across the entire cohort, expression was not significantly different between genotypes (unpaired t-test with Welch’s correction, p=0.8584). However, as shown in **Figure 1C**, we observed that three samples that were separated in expression from the remainder of the RTT group via a ROUT test; when these samples were removed and the expression of *HRH3* of the remaining RTT samples was compared to control, we found a significant decrease in the remaining RTT samples (**Figure 1C**).

**Figure 1.**
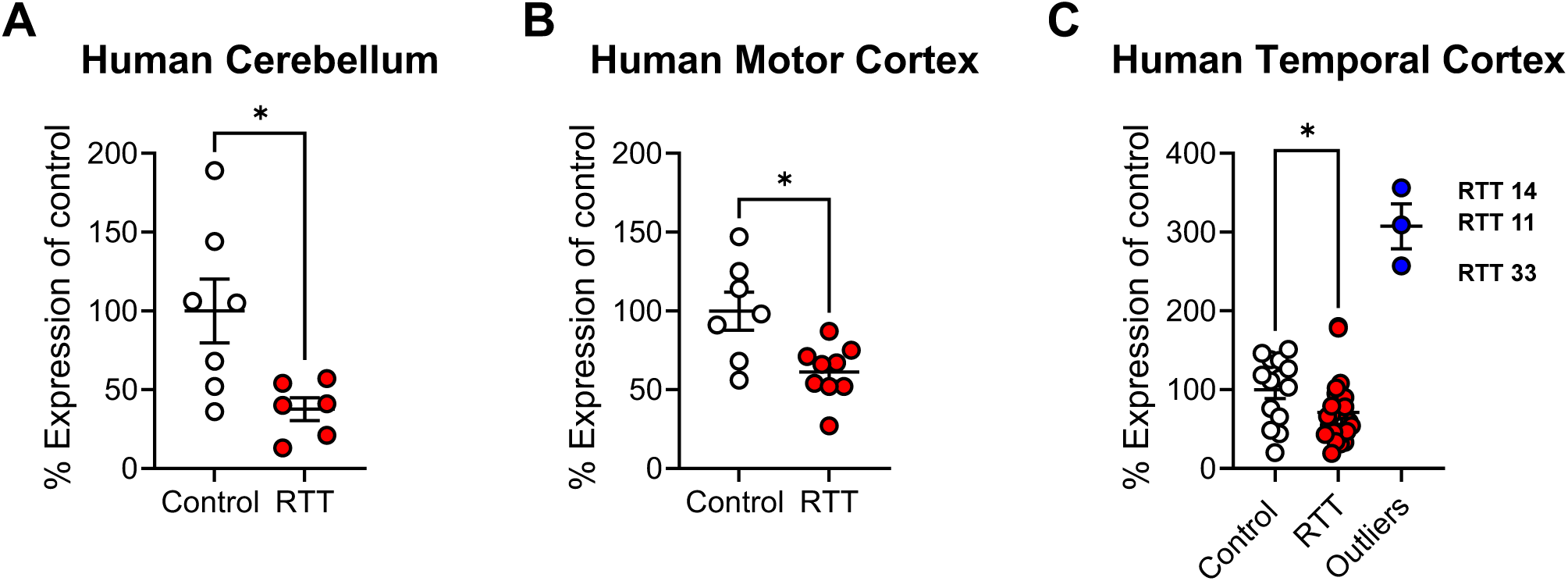
Histamine H_3_ receptor (*HRH3*) expression is reduced in the brains of RTT patients with defined *MECP2* mutations. Quantitative RT-PCR was performed and expression was normalized to the control group. **A.** *HRH3* expression in the original cohort in the cerebellum of control (n=7, white) and RTT (n=6, red) patients was significantly decreased (Mean ± SEM; unpaired t-test with Welch’s correction, *p = 0.0205, t(7.449) = 2.892). **B.** *HRH3* expression in the original cohort in the motor cortex of control (n=7, white) and RTT (n=9, red) patients was significantly decreased (Mean ± SEM; unpaired t-test with Welch’s correction, *p = 0.0183, t(8.736) = 2.896). **C.** Profiling of *HRH3* expression in a larger cohort from temporal cortex from patients with identified *MECP2* mutations (Table 1) revealed that the majority of RTT patients (n=25, red) exhibited significantly decreased *HRH3* levels compared to controls (n=14, white, Mean ± SEM, unpaired t-test with Welch’s correction between control and RTT, *p = 0.0447, t(25.85) = 2.109) across the cohort, with the exception of three patients (blue).

The observation that there were reductions in *HRH3* expression in the brains of many of the patients replicated the work of Renthal et al. from the occipital cortex (Renthal, Boxer et al. 2018), suggesting that the H_3_ receptor might serve as a novel, druggable therapeutic target in RTT. To translate our findings to animals, we next evaluated mRNA expression of the mouse H_3_ receptor in samples from *Mecp2^Null/+^* female animals via qRT-PCR in brain samples from the cortex, hippocampus, striatum and cerebellum (**Figure 2A-D**). These studies revealed significant decreases in *Hrh3* receptor mRNA expression in the cortex and striatum in 20-week-old *Mecp2^Null/+^* animals compared to wild-type controls.

**Figure 2.**
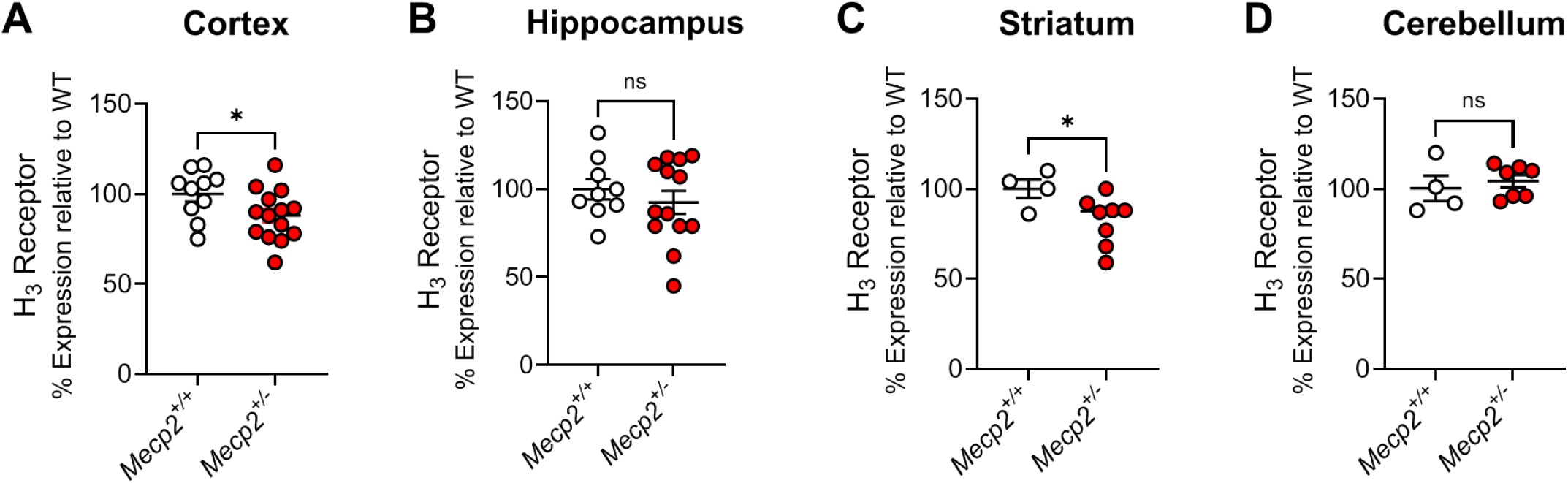
Mouse *Hrh3* mRNA expression is decreased in the cortex and striatum of *Mecp2^+/-^* mice compared to controls. Quantitative RT-PCR was performed on samples from various brain areas for *Mecp2^+/+^* (white) and *Mecp2^Null/+^* (red) mice. Comparisons were unpaired Student’s t-tests; cortex, t(22) = 2.101; *p = 0.0474; hippocampus, t(20) = 0.8123, p = 0.4262; striatum, t(10) = 2.288, *p = 0.0451; cerebellum, t(9) = 0.5872, p = 0.5715.

The H_3_ receptor agonist, RAMH, does not affect locomotion in an open field but increases time in the open arms in an elevated zero maze in *Mecp2^Null/+^* mice. We evaluated the effects of the compound *(R)*-α*-*methylhistamine (RAMH, 45 mg/kg, ip), an agonist of the H_3_ receptor (Arrang, Devaux et al. 1988, Arrang, Garbarg et al. 1988, Rapanelli, Frick et al. 2016) on several behavioral phenotypes in symptomatic 20-week-old *Mecp2^+/+^* and *Mecp2^Null/+^* female RTT model mice: locomotion in an open field, anxiety as assessed by elevated zero maze, repetitive behavior using a hindlimb clasping measurement, and learning and memory in a Pavlovian task, conditioned fear. In the open field assessment, mice were administered either RAMH or vehicle 15 minutes prior to testing and then monitored in the open field for 60 minutes (**Figure 3A**). Vehicle-treated *Mecp2^Null/+^* animals exhibited reductions in distance traveled in the early time points of the task at 5 and 10 minutes compared to *Mecp2^+/+^* controls (**Figure 3A**), but this effect between vehicle-treated animals was not significant once ambulation was calculated across the entire 60-minute evaluation period (**Figure 3B**). *Mecp2^Null/+^* treated with RAMH also did not exhibit significantly different total distance traveled compared to vehicle-treated *Mecp2^Null/+^* mice (**Figure 3B**). In *Mecp2^+/+^* controls, RAMH also had no effect. If the open field arena was parsed into the duration in the center versus the surround, there were no differences between genotypes and treatments (**Figure 3C, D**).

**Figure 3.**
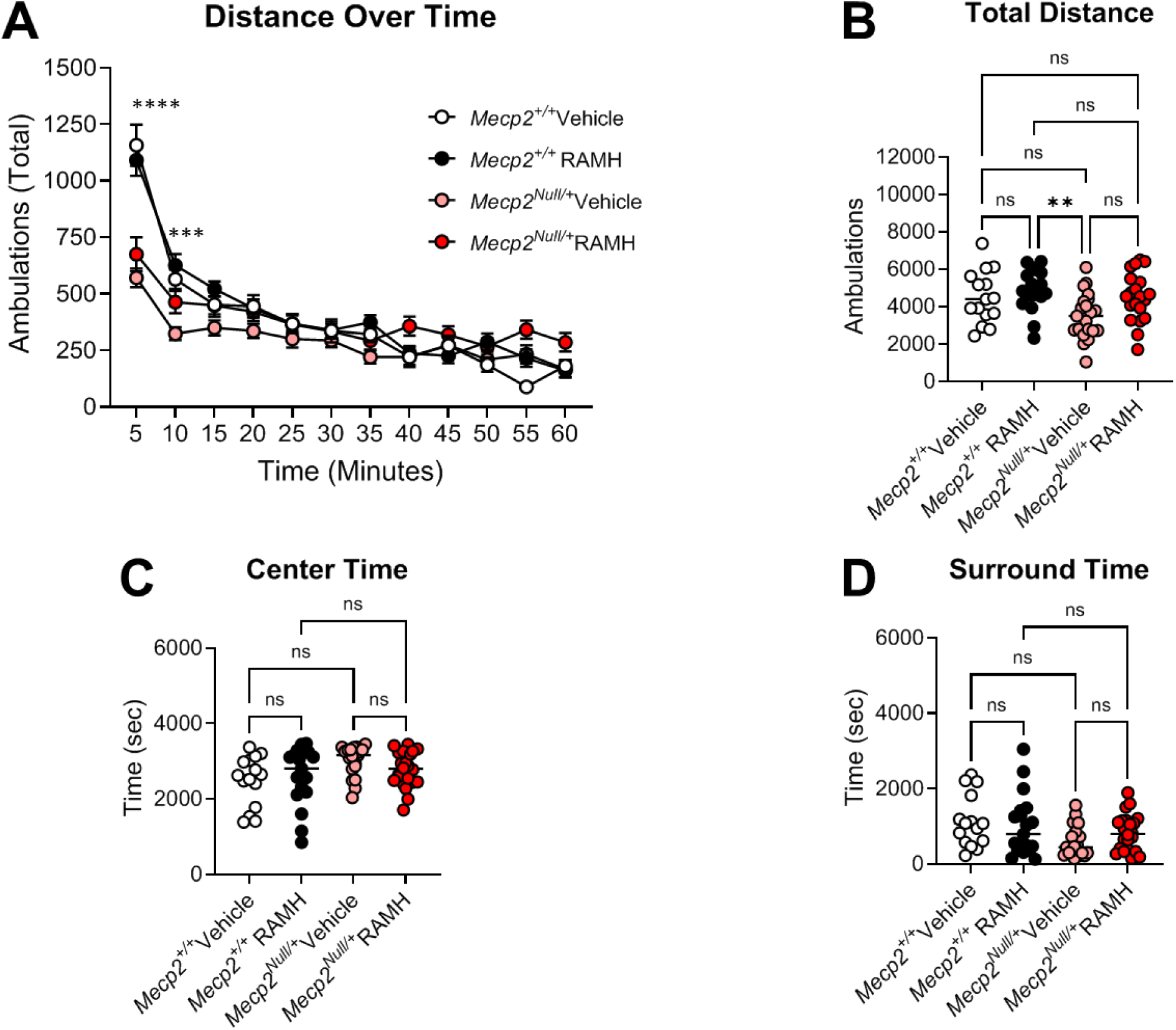
The H_3_ agonist RAMH does not affect locomotion in *Mecp2^+/-^* mice. *Mecp2^+/+^* and *Mecp2^Null/+^* mice were administered vehicle or 45 mg/kg of RAMH 15 minutes prior to placement in an open field and allowed to explore the area for 60 minutes. **A.** Time course of exploration. *Mecp2^+/+^* vehicle, white symbols; *Mecp2^+/+^* RAMH, black symbols; *Mecp2^Null/+^* vehicle, pink symbols; *Mecp2^Null/+^* RAMH, red symbols. 2-way ANOVA revealed significant differences in interaction between groups and time (F (18.16, 472.1) = 8.915, ****p < 0.0001) and genotype and treatment (F (3, 78) = 6.730, ***p = 0.0004).Tukey’s post hoc test revealed differences between *Mecp2^+/+^* vehicle and *Mecp2^Null/+^* vehicle at timepoints 5 (****p < 0.0001) and 10 (***p < 0.001) minutes. **B.** When quantified over 60 minutes (Ordinary 1-way ANOVA, F (3, 71) = 4.508, **p = 0.0060), RAMH-treated *Mecp2^+/+^* mice and vehicle-treated *Mecp2^Null/+^* mice were significantly different (Tukey’s post hoc test between vehicle-treated *Mecp2^+/+^* RAMH mice and vehicle-treated *Mecp2^Null/+,^* **p =0.0050. No other effects between genotypes or treatment were apparent). **C.** When time in the center was measured, there was no significant effect of genotype or treatment (Ordinary 1-way ANOVA, F (3, 71) = 2.616, p = 0.0577). **D.** When time in the surround was measured, no difference in time was observed for either genotype or treatment (Ordinary 1-way ANOVA, F (3, 71) = 2.660, p = 0.0547).

To assess anxiety, we performed an elevated zero maze assessment (EZM; **Figure 4A**), and mice were again administered RAMH or vehicle 15 minutes prior to testing. In this assay, increased time spent in the open arms of the maze is interpreted as anxiolytic, and *Mecp2^Null/+^* treated with RAMH exhibited significantly higher amounts of time in the open arms compared to *Mecp2^Null/+^* mice treated with vehicle (**Figure 4A**). This effect was not present in the *Mecp2^+/+^* animals. We also observed no differences in the number of entries into the open arms or the distance traveled in this maze task based on genotype or treatment (**Figure 4B and C**) or distance in the open arms of the maze.

**Figure 4.**
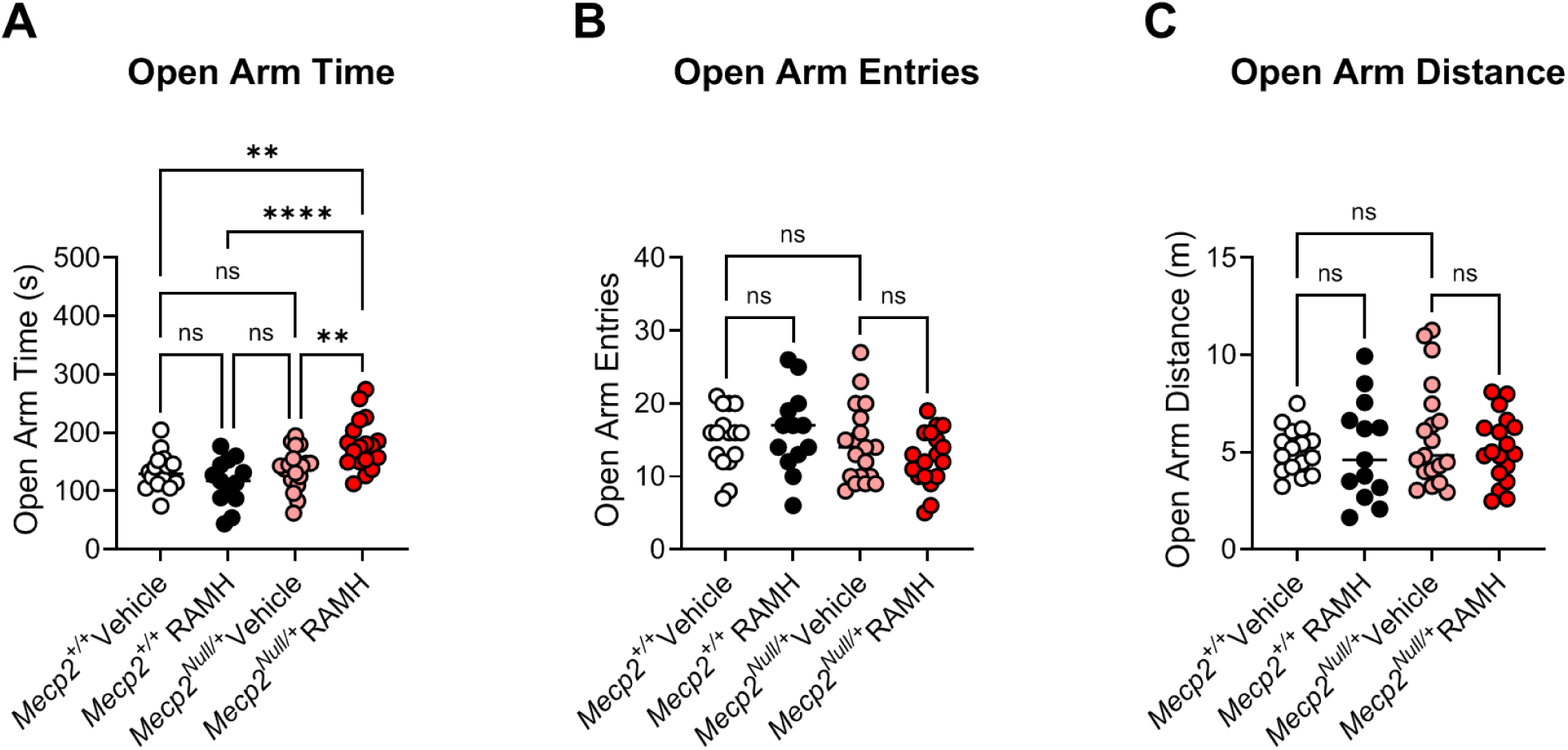
RAMH increases the amount of time *Mecp2^Null/+^* mice spend in the open arms of the elevated zero maze. **A.** When time in the open arms of an elevated zero maze is measured, *Mecp2^Null/+^* mice treated with RAMH spend significantly more time in the open arms compared to vehicle-treated *Mecp2^Null/+^* animals or *Mecp2^+/+^* controls (Ordinary 1-way ANOVA, F (3, 62) = 8.740, ****p <0.0001; Tukey’s post hoc test, **p = 0.0028 between *Mecp2^+/+^* vehicle (white) and *Mecp2^Null/+^* RAMH (red); ****p < 0.0001, *Mecp2^+/+^* RAMH (black) versus *Mecp2^Null/+^* RAMH (red); **p = 0.0057 between *Mecp2^Null/+^* vehicle (pink) and *Mecp2^Null/+^* RAMH (red)). **B and C.** There was no significant change in number of entries (1-way ANOVA, F (3, 62), 1.709, p = 0.1743) or distance (1-way ANOVA, F (3, 62), 0.6073, p = 0.6127) in the open arms of the maze.

Patients with RTT exhibit a repetitive hand wringing phenotype which prevents purposeful hand movements and usage (Weng, Bailey et al. 2011, Townend, Ehrhart et al. 2018, Stallworth, Dy et al. 2019). *Mecp^Null/+^* mice exhibit a clasping phenotype of their hindlimbs (Guy, Hendrich et al. 2001, Guy, Gan et al. 2007). To assess the effects of H_3_ activation in clasping, mice were administered RAMH or vehicle 15 minutes prior to an assessment of hindlimb clasping over one minute. This evaluation showed the expected clasping phenotype of the *Mecp2^Null/+^* mice versus controls in the presence of vehicle (***p < 0.0001), but there was no effect of RAMH on this phenotype (**Figure 5A**). Finally, we assessed activity in a contextual fear conditioning task; we did not observe impairments in contextual fear conditioning in RTT mice in this cohort and administration of RAMH did not affect the performance of mice in this task (**Figure 5B**).

**Figure 5.**
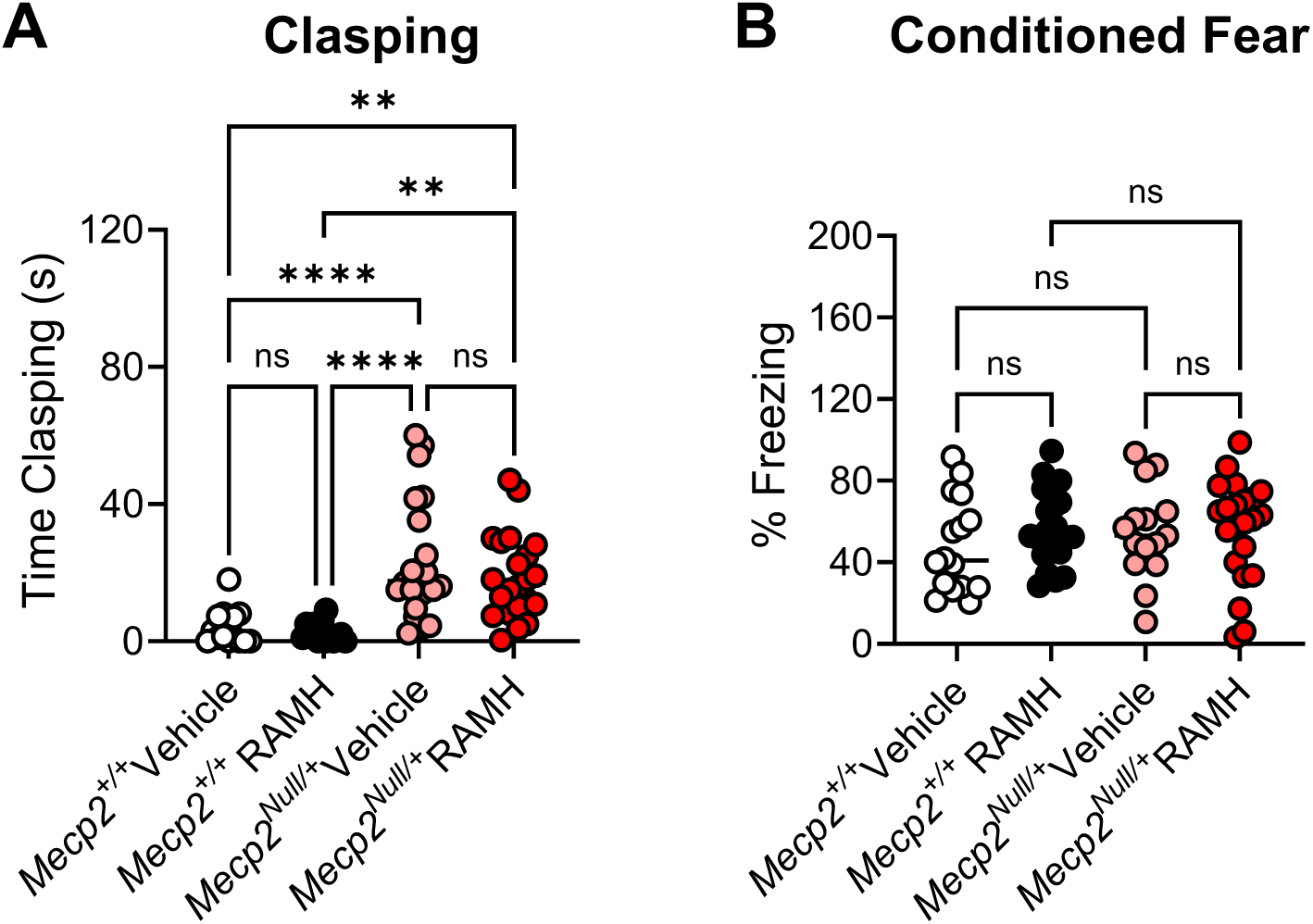
RAMH does not affect hindlimb clasping or contextual fear conditioning responses in *Mecp2^Null/+^* animals. **A.** Hindlimb clasping was assessed for one minute by suspending animals from their tail. *Mecp2^Null/+^* animals treated with vehicle exhibited significantly higher levels of clasping compared with littermate *Mecp2^+/+^* controls, and this was not corrected with the administration of RAMH. Ordinary 1-way ANOVA, F (3, 69), 13.20, ****p < 0.0001, followed by Tukey’s post-hoc test: *Mecp2^+/+^* Vehicle (white) versus *Mecp2^Null/+^* Vehicle (pink); ****p < 0.0001; *Mecp2^+/+^* Vehicle (white) versus *Mecp2^Null/+^* RAMH (red), **p = 0.0029; *Mecp2^+/+^* Vehicle (white) versus *Mecp2^+/+^* RAMH, p = 0.9865; *Mecp2^+/+^* RAMH (black) versus *Mecp2^Null/+^* Vehicle (pink), ****p < 0.0001; *Mecp2^+/+^* RAMH (black) versus *Mecp2^Null/+^* RAMH (red), **p = 0.0011; *Mecp2^Null/+^* Vehicle (pink), versus *Mecp2^+/+^* RAMH (red), p=0.5763. **B.** No differences were noted in contextual fear conditioning responses (Ordinary 1-way ANOVA, F (3, 71), 0.6006, p = 0.6167).

### A mouse model of *MECP2* duplication syndrome (MDS) expresses elevated histamine H_3_ mRNA receptor expression in cortex and striatum

The decreased expression of the histamine H_3_ receptor in RTT patients and the *Mecp2^Null/+^* mouse model of RTT, as well as previous literature, prompted us to examine *Hrh3* expression in *MECP2^Tg1^* animals, a model of MDS (Collins, Levenson et al. 2004). We quantified mRNA expression in cortex, hippocampus, striatum, and cerebellum of male and female *MECP2^Tg1^* animals (**Figure 6**). These studies showed that there was a significant increase in *Hrh3* expression in the cortex and striatum of *MECP2^Tg1^* animals compared to wild-type (WT) littermate controls, but not in the hippocampus or cerebellum. The compound pitolisant, a histamine H_3_ receptor antagonist, has recently been approved for use in humans to treat sleep disorders, specifically narcolepsy (Wakix®, (Fabara, Ortiz et al. 2021, Sarfraz, Okuampa et al. 2022)) and has been shown to exhibit efficacy in fear memory in wild-type mice (Brabant, Charlier et al. 2013). This suggested that pitolisant might have efficacy in normalizing abnormal behavioral phenotypes, particularly those requiring intact cortical and striatal circuitry, in *MECP2^Tg1^* animals. We performed a similar battery of studies in *MECP2^Tg1^* mice and WT controls, progressing animals through open field, EZM, and conditioned fear (*MECP2^Tg1^* mice do not exhibit a clasping phenotype); pitolisant (10 mg/kg, ip) was administered 30 min prior in these studies.

**Figure 6.**
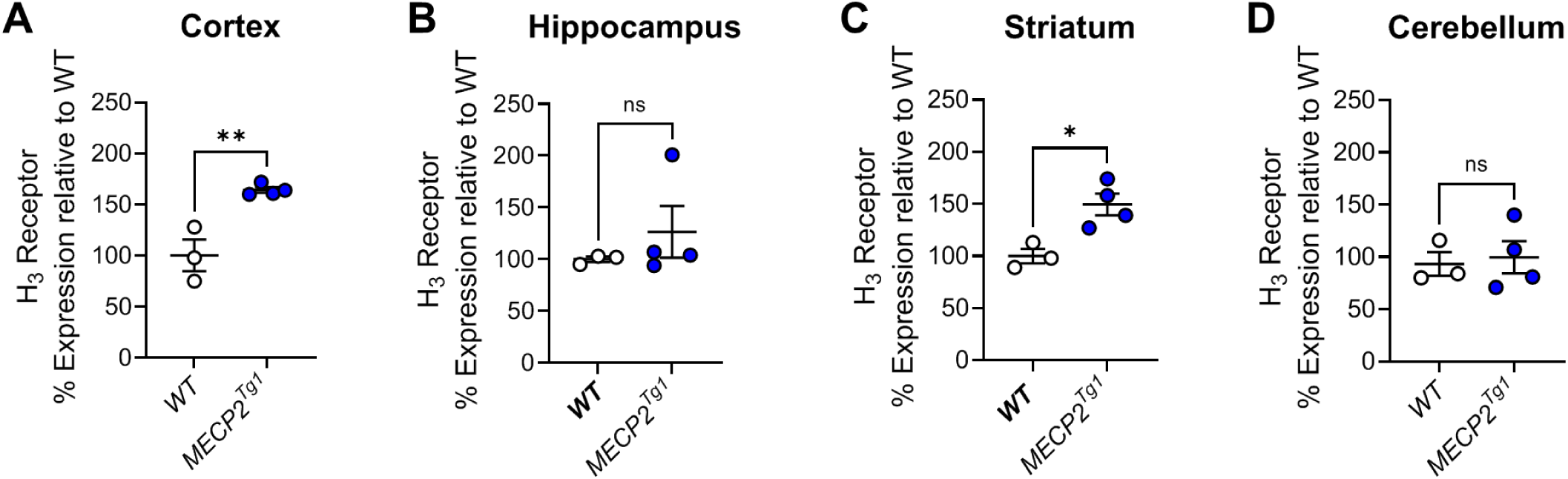
Mouse *Hrh3* expression is increased in the cortex and striatum of *MECP2^Tg1^* mice compared to controls. Quantitative RT-PCR was performed on samples from various brain areas for wild-type littermate controls (WT, white) or *MECP2^Tg1^* (blue) mice. Statistical comparisons were unpaired t-tests; cortex, t(5) = 4.829, **p = 0.0048; hippocampus, t(5) = 0.8940, p = 0.4123; striatum, t(5) = 3.642, *p = 0.0149; cerebellum, t(5) = 0.3118, p = 0.7678; n=3-4 samples per tissue.

### Pitolisant does not correct behavioral deficits in *MECP2^Tg1^* animals

In an open field test, we observed no differences in total distance traveled when sexes were combined (**Figure 7A, B**). Separation into male and female animals revealed that, while no significant differences were identified in the males (**Supplemental Figure 1A**), *MECP2^Tg1^* females exhibited reduced distance traveled compared to WT females (**Supplemental Figure 1B**). Additionally, WT females, but not *MECP2^Tg1^* females, exhibited a significant reduction in locomotion in response to pitolisant (**Supplemental Figure 1B**). There were no differences in time spent in the center or the surrounding area of the open field (**Figure 7C, D, Supplemental Figure 1C-F**). In EZM, we observed a reduction in the amount of time spent in the open arms in *MECP2^Tg1^* animals compared to WT controls; however, this phenotype was not altered by pitolisant (**Figure 8A**). When mice were separated by sex, there was a significant difference in open arm time only in the male animals (*MECP2^Tg1^* versus WT, **Supplemental Figure 1G, H**), but, again, no effects of pitolisant were observed. Additionally, there were no significant differences in open arm entries or open arm distance traveled (**Figure 8B**, **C, Supplemental Figure 1I-L**). Finally, similar to the findings of multiple labs, *MECP2^Tg1^* mice treated with vehicle exhibited a strong response in conditioned fear in which they froze significantly more than WT animals ((Na, Nelson et al. 2012, Stansley, Fisher et al. 2018, Vermudez, Buch et al. 2022), **Figure 9)**. This was most prominently observed in female animals (**Supplemental Figure 1M-N**); however, this phenotype was not impacted by pitolisant in either sex, suggesting that pitolisant does not correct the behavioral deficits tested here in *MECP2^Tg1^* animals

**Figure 7.**
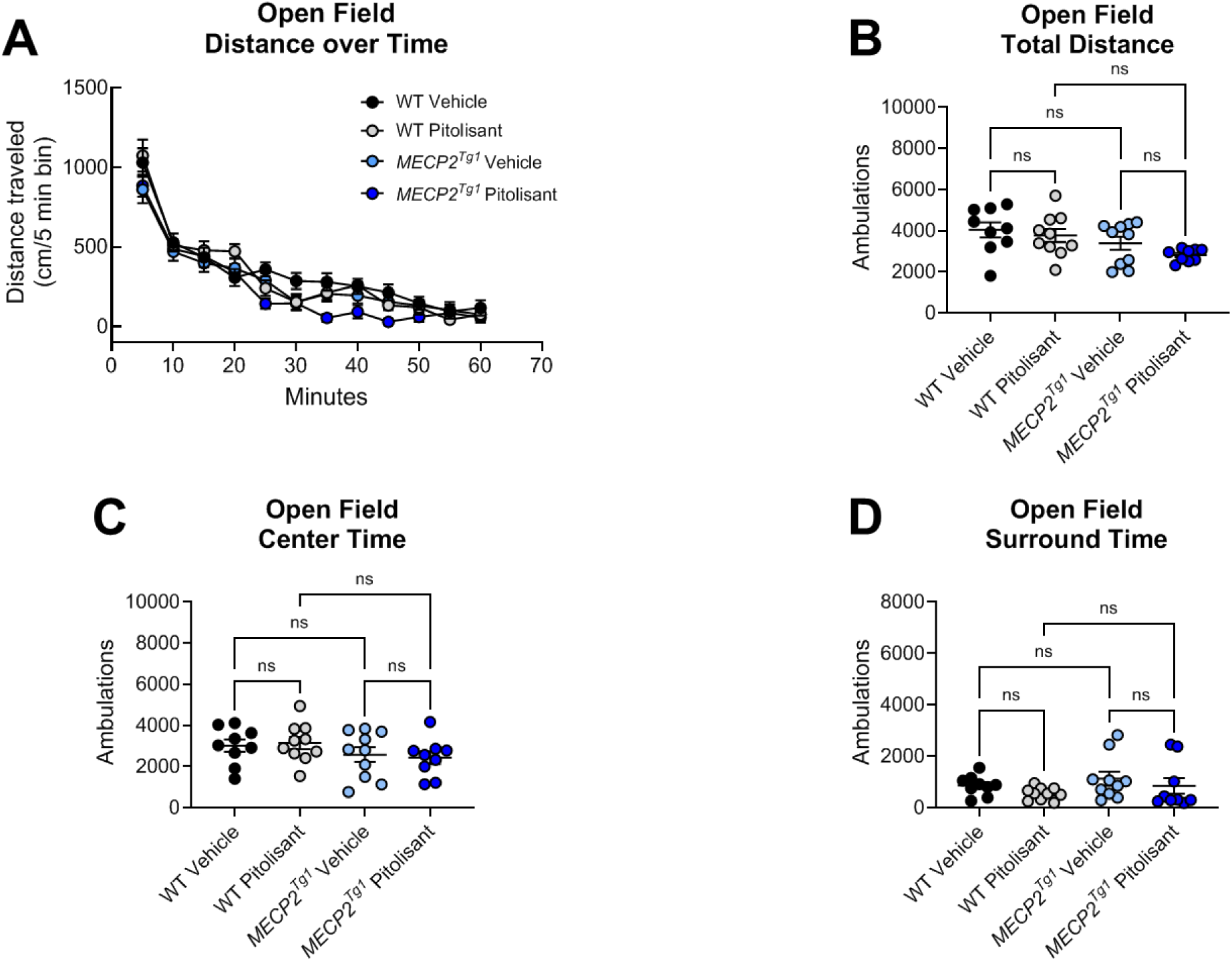
The H_3_ receptor antagonist pitolisant does not impact open field behavior in *Mecp2^+/+^*or *MECP2^Tg1^* mice. Wild-type (WT) littermate controls and *MECP2^Tg1^* mice were administered vehicle or 10 mg/kg of pitolisant 30 minutes prior to placement in an open field and allowed to explore the area for 60 minutes. **A.** Time course of exploration. WT vehicle, black symbols; WT pitolisant, gray symbols; *MECP2^Tg1^* vehicle light blue symbols; *MECP2^Tg1^* pitolisant, dark blue symbols. **B.** Total distance traveled over 60 minutes (Ordinary 1-way ANOVA, F (3,34), 2.978, p = 0.0451; Tukey’s post-test, WT vehicle and *MECP2^Tg1^* pitolisant, *p=0.0393; all other comparisons were not significant). **C.** Distance in the center was not different between genotypes or treatments (Ordinary 1-way ANOVA, F (3, 34), 1.149, p = 0.3434). **D.** Distance in the surrounding area was not different between genotypes or treatments (1-way ANOVA, F (3, 34), 1.149, p = 0.3434).

**Figure 8.**
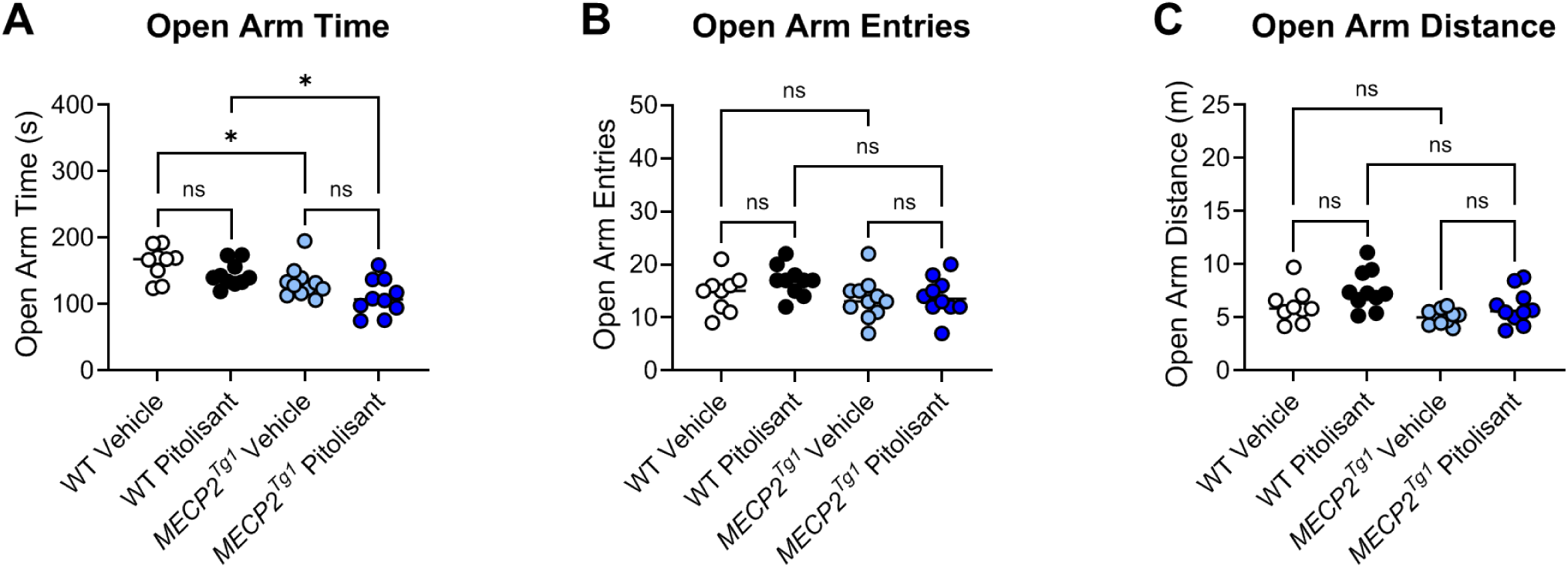
Pitolisant does not exhibit efficacy in elevated zero maze. **A.** *MECP2^Tg1^* mice exhibited a significantly decreased in time in the open arms of an elevated zero maze compared to wild-type (WT) littermate controls, but this response was not affected by pitolisant (Ordinary 1-way ANOVA, F (3, 36), 7.834, ***p = 0.0004; Tukey’s post hoc test, WT vehicle (white) versus *MECP2^Tg1^* vehicle (light blue), *p = 0.0447; WT vehicle (white) versus *MECP2^Tg1^* pitolisant (dark blue), ***p = 0.0002; WT pitolisant (black) versus *MECP2^Tg1^* pitolisant (dark blue), *p = 0.0162; all other comparisons were not significant). **B.** There was no significant change in number of entries (Ordinary 1-way ANOVA, F (3, 36), 1.958, p = 0.1378). **C.** The only significant open arm distance comparison was WT pitolisant (black) versus *MECP2^Tg1^* vehicle (light blue) (not shown on graph, 1-way ANOVA, F (3, 35), 4.764, **p = 0.0069); Tukey’s post hoc test, **p = 0.0035); all other comparisons were not significant.

**Figure 9.**
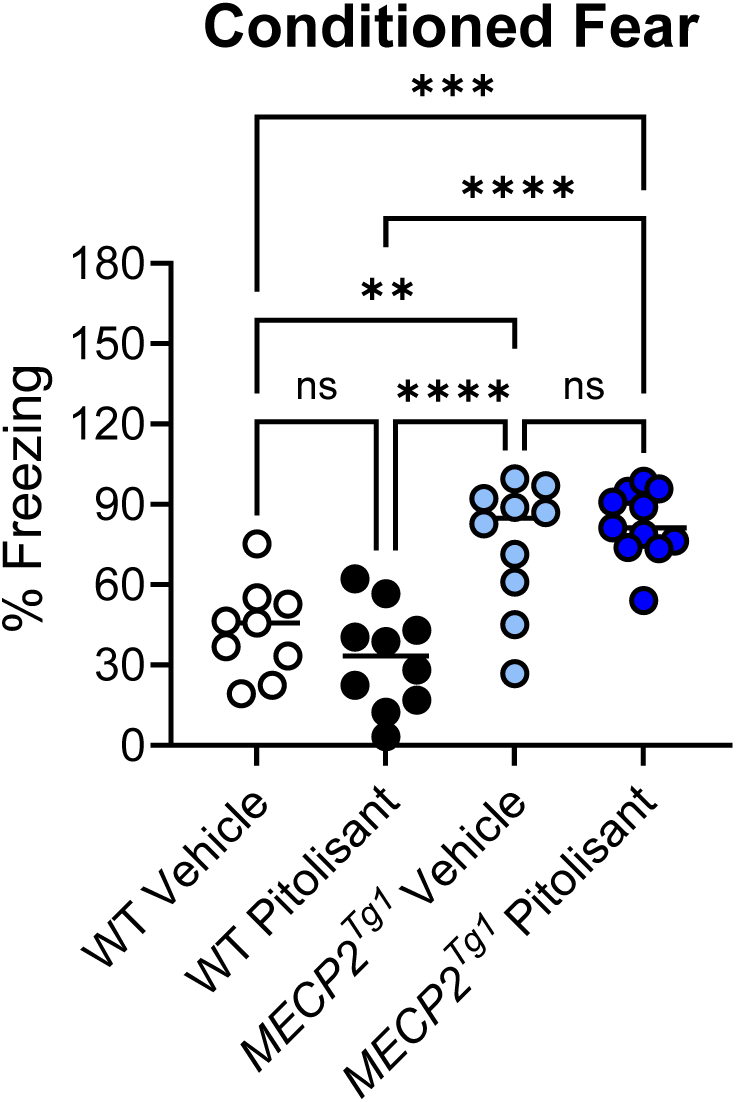
Pitolisant does not change the elevated contextual fear response characteristic of *MECP2^Tg1^*animals. Wild-type (WT) littermate controls and *MECP2^Tg1^* mice were treated with vehicle or pitolisant 30 minutes before training in a contextual fear conditioning task; percent freezing was assessed 24 hours later when the mice were returned to the same context (Ordinary 1-way ANOVA, F (3.36), 17.09, ****p< 0.0001).Tukey’s post-test, WT Vehicle versus WT pitolisant, p = 0.6119; WT Vehicle versus *MECP2^Tg1^* Vehicle, **p = 0.0035; WT Vehicle versus *MECP2^Tg1^* pitolisant, ***p = 0.0002; WT pitolisant versus *MECP2^Tg1^* Vehicle, ****p < 0.0001; WT pitolisant versus *MECP2^Tg1^* pitolisant, ****p < 0.0001; *MECP2^Tg1^* Vehicle versus *MECP2^Tg1^* pitolisant, p = 0.8106.

## Discussion

RTT is a disorder characterized by a constellation of symptoms that appear in girls following a regression period, including the development of intellectual disability, seizures, repetitive behaviors, and anxiety. MDS is a related disorder that results in many of the same symptom domains, often with opposing phenotypes. We and others have taken a “bedside-to-bench” approach to identify targets with clinically relevant changes in expression using ‘omics profiling of samples from patients and model mice. These strategies have resulted in our exploration of the therapeutic potential of several mGlu and muscarinic targets (Gogliotti, Senter et al. 2016, Gogliotti, Senter et al. 2017, Gogliotti, Fisher et al. 2018, Cikowski, Holt et al. 2022, Smith, Arthur et al. 2022, Vermudez, Buch et al. 2022). We have now turned our attention to the H_3_ histamine receptor based on findings that expression of the receptor is reduced in autopsy samples from a cohort of RTT patients, correlating with previous data published by (Renthal, Boxer et al. 2018), and we validated this decrease in expression across a larger cohort of RTT samples. These findings reveal that the majority of RTT patients can be clustered into a group with significantly reduced *HRH3* expression (**Figure 1C**). Interestingly, there appear to be several individuals who diverge from this pattern; with the exception of potentially the P302L/R306C samples (2 patients expressed *HRH3* levels at 309 and 356% of control, one patient expressed 101% of control), this does not appear to correlate with specific *MECP2* mutation but we are limited due to the low number of patients with each mutation.

The *Mecp2^Null/+^* mouse model of RTT provides strong face and construct validity for the study of potential treatments (Guy, Hendrich et al. 2001, Vashi and Justice 2019). These mice exhibit numerous behavioral phenotypes that are similar to those observed in the human disorder, notably, impairments in cognition, motor impairments, seizures, repetitive behaviors, and anxiety (Guy, Hendrich et al. 2001). Additionally, replacement of Mecp2 corrects behavioral deficits and restores normal lifespan (Guy, Gan et al. 2007) suggesting that the disease is not degenerative in mice. Therapeutic strategies for RTT have now taken several avenues: those aimed at correcting MeCP2 itself using strategies such as gene therapy, DNA and RNA editing, and X chromosome reactivation, and strategies that exploit changes in targets that occur upon the loss of MeCP2. While only the former strategies would be considered curative, symptomatic treatments may still be of benefit in the daily lives of RTT patients. Furthermore, the overlap of symptoms with other neurodevelopmental disorders suggests that treatments that impact RTT phenotypes may extend to other disorders. After determining that there are reduced levels of *HRH3* mRNA in postmortem brain samples of RTT patients, we also found here that 20-week-old, symptomatic *Mecp2^Null/+^* mice express reduced levels of *Hrh3* mRNA in the cortex and striatum when compared to *Mecp2^+/+^* controls, but normal levels in the hippocampus and cerebellum. The finding that *Hrh3* mRNA level is not reduced in the cerebellum of *Mecp2^Null/+^* mice compared to *Mecp2^+/+^* controls contrasts with our original RNA sequencing data from human RTT patients, where we observed decreased expression of *HRH3* in the cerebellum. This discrepancy might be attributed to species-specific differences in the expression and distribution of histamine H_3_ receptor mRNA in the cerebellum (Li, Zhu et al. 2014). Indeed, we and others have previously observed low overlap of gene expression changes between mouse models and RTT patients (Gogliotti, Fisher et al. 2018, Renthal, Boxer et al. 2018). We would also note that our findings of reductions in *HRH3* mRNA in the RTT cortex align with those previously reported in other RNA sequencing studies (Ben-Shachar, Chahrour et al. 2009, Chao, Chen et al. 2010)

RTT patients exhibit a variety of repetitive behaviors and anxiety phenotypes (Stallworth, Dy et al. 2019, Buchanan, Stallworth et al. 2022). These include the repetitive hand wringing/clasping that is a hallmark of the disorder and a core diagnostic criterion for RTT (Gold, Percy et al. 2024). Anxiety symptoms range from self-injurious behavior, screaming and crying, and mood changes that are common among RTT patients (Barnes, Coughlin et al. 2015, Gold, Percy et al. 2024). In mouse models of RTT, specific behavioral phenotypes can be separated when MeCP2 is lost from or replaced within specific neuronal subtypes (Chao, Chen et al. 2010, Ito-Ishida, Ure et al. 2015, Meng, Wang et al. 2016, Ure, Lu et al. 2016). Interestingly, histamine H_3_ receptors have been shown to be involved in the manifestation of anxiety behaviors (Yokoyama, Yamauchi et al. 2009, Mohsen, Yoshikawa et al. 2014, Zhang, Peng et al. 2020). Mechanistic studies revealed that activation of the *Hrh3* system in the nucleus accumbens (NAc) core inhibits glutamatergic neurotransmission from prelimbic regions to the NAc circuit and inhibits anxiety-like and obsessive-compulsive-like behavior induced by exposure to stress (Zhang, Peng et al. 2020). These studies indicate that the histamine H_3_ receptor might be a target for developing potential therapeutic strategies for anxiety and repetitive behaviors. Additionally, mice fed a low histidine diet exhibit reductions in histamine levels and release in the brain (Yoshikawa, Nakamura et al. 2014) and develop anxiety-relevant behaviors. RTT patients have also been shown to have lower levels of histidine in plasma (Neul, Skinner et al. 2020) and experience multiple sleep-related disorders (Tascini, Dell’Isola et al. 2022). Together, these results suggest a strong link between H_3_ expression and histamine levels with anxiety-like phenotype in RTT model mice, which provided a rationale to test 20-week-old *Mecp2^Null/+^* mice in a behavior relevant to anxiety. While much attention has been focused on the development of antagonists/inverse agonists for the H_3_ receptor (Fabara, Ortiz et al. 2021, Keam 2023), several compounds that increase the activity of H_3_ receptors have also been tested clinically for indications such as asthma and migraine (O’Connor, Lecomte et al. 1993, Fozard 2000, Leurs, Bakker et al. 2005, Millan-Guerrero, Isais-Millan et al. 2006). Using a pharmacological treatment scheme comparing the effects of 45 mg/kg of the histamine H_3_ receptor agonist, RAMH, significantly increased the time spent by *Mecp2^Null/+^* mice in open arms of the elevated zero maze, suggesting that increasing H_3_ receptor signaling may alleviate anxiety-like behaviors. This effect was not accompanied by significant reductions in arm entries or in total distance traveled, suggesting that the effects on open arm time are not confounded by decreases in movement. It should be noted that effects in the open field in various lines of RTT mice in general are conflicting, with some reports showing that RTT model mice spend less time in the open arms of the maze (interpreted as higher anxiety) and other studies showing that RTT mice spend more time in the open arms (interpreted as lower anxiety) at baseline (Gogliotti, Senter et al. 2017). Here, we did not observe a significant difference in open arm time between *Mecp2^+/+^* and *Mecp2^Null/+^* mice with vehicle treatment, but, as a group, *Mecp2^Null/+^* animals treated with RAMH exhibited a robust preference for the open arms of the maze. This suggests a clear effect of RAMH that is only present in the *Mecp2^Null/+^* animals.

Activation of H_3_ has been shown to ameliorate obsessive-compulsive-like behaviors induced by activation of cortico-accumbal glutamatergic afferents (Zhang, Peng et al. 2020). Based on these findings and due to the robust hindlimb clasping phenotype observed in *Mecp2^Null/+^* mice (Patterson, Hawkins et al. 2016), we anticipated that this might be a domain in which *Mecp2^Null/+^* females might be sensitive to RAMH treatment. In contrast to this hypothesis, we did not observe efficacy of RAMH in reversing hindlimb clasping. This observation fits with the findings of the Zoghbi lab in which anxiety and hindlimb clasping were separated when Mecp2 was restored in glutamatergic versus GABAergic neurons in mice, respectively (Meng, Wang et al. 2016, Ure, Lu et al. 2016). Future studies in this area could include testing the efficacy of RAMH in mice with loss of Mecp2 selectively from each type of neuron; if the above hypothesis is correct, it would be anticipated that the effects of RAMH on anxiety phenotypes would be lost in the mice lacking Mecp2 in glutamatergic, but not GABAergic, neurons. Indeed, RAMH-induced activation of presynaptic H_3_ receptors has been shown to selectively inhibit glutamatergic, but not GABAergic, neurotransmission in the NAc core and activation of these receptors alleviates anxiety-like behaviors induced by optogenetic activation of glutamatergic signaling from the prelimbic cortex to the NAc core circuitry (Zhang, Peng et al. 2020). Since *Mecp2* is also expressed in cortical areas and in the NAc, studies aiming at determining whether loss of Mecp2 expressed in this circuitry mediates anxiety-like behavior observed in *Mecp2^Null/+^* mice is an exciting future direction.

Tight regulation of MeCP2 expression is important as duplication of the gene causes MDS with considerable overlap with classical RTT syndrome patients in terms of the presence of stereotypic behavior, anxiety, and social avoidance (Moretti and Zoghbi 2006, Ramocki, Peters et al. 2009). Therefore, similar to our studies in *Mecp2^Null/+^* mice, we also profiled the levels of histamine H_3_ receptors in the cortex, hippocampus, striatum and cerebellum of *MECP2^Tg1^* mice. In contrast to the *Mecp2^Null/+^* animals, we observed an elevated expression of histamine H_3_ receptors mRNA in the cortex and striatum of *MECP2^Tg1^* mice, suggesting elevated histamine H_3_ receptor signaling in these areas. This finding prompted us to test the activity of the histamine H_3_ receptor antagonist, pitolisant (Fabara, Ortiz et al. 2021, Sarfraz, Okuampa et al. 2022), in correcting observed behavioral deficits in a mouse model of MDS. In contrast to our hypothesis, pitolisant did not correct anxiety-like behavior and enhanced freezing behavior in *MECP2^Tg1^* mouse model of MDS, suggesting that inhibition of histamine H_3_ receptor signaling with pitolisant was not sufficient to correct these behaviors in MDS mice. It should be noted that the main indication for pitolisant in the clinic is narcolepsy and sleep disorders, which were not examined here. Patients with MDS have been reported to exhibit sleep disturbances (Pehlivan, Huang et al. 2025) and 80% of patients with RTT exhibit sleep problems (Tascini, Dell’Isola et al. 2022, Gold, Percy et al. 2024). Future studies examining H_3_ receptor modulation in sleep in RTT and MDS mice, therefore, may be areas that deserve future study.

In summary, our findings reveal that administration of histamine H_3_ receptor agonist, RAMH, can increase time spent by *Mecp2^Null/+^* mice in the open arms of an elevated zero maze test of anxiety. Future studies designed to evaluate the performance of mice in a battery of anxiety tests like elevated plus maze and light/dark transition task will provide more information about the role of histamine H_3_ receptors in regulating anxiety in RTT mice, as these test measure anxiety under different conditions (van Gaalen and Steckler 2000). Additionally, future studies could explore the effect of chronic dosing of an H_3_ agonist. Overall, these studies provide important information regarding the physiological role of histamine H_3_ receptor in the regulation of anxiety behavior in RTT syndrome.

## Supporting information

Supplemental Fig. 1. Pitolisant effects in male versus female MECP2Tg1 animals and wild-type (WT) littermate controls.

## Authorship contribution statement

**Kelly Weiss:** Writing – original draft, Visualization, Methodology, Investigation. **Sheryl Anne D. Vermudez:** Writing – original draft, Visualization, Methodology, Investigation. **Geanne A. Freitas:** Writing – review & editing, Visualization, Methodology, Investigation. **Mackenzie Meadows:** Visualization, Methodology, Investigation. **Shalini Dogra:** Writing – review & editing, Visualization. **Rocco Gogliotti:** Writing – review and editing, Visualization, Methodology, Investigation, Supervision, Resources, Project administration, Methodology, Funding acquisition, Formal analysis, Conceptualization. **Colleen M. Niswender:** Writing – review & editing, Writing – original draft, Supervision, Resources, Project administration, Methodology, Funding acquisition, Formal analysis, Conceptualization.

## Data availability

All mentioned data are presented in this published article or the supplementary information or are available from the corresponding author upon reasonable request.

## Ethics statement

This manuscript is an original work and is not under consideration for publication elsewhere. All authors have reviewed and approved of the submission.

## Funding

This work was supported by grants R01MH124671, R01NS031373, R21MH102548, R01MH113543, K01MH112983, R01NS112171, and the International Rett Syndrome Foundation. Harvard Brain Tissue Resource Center is supported by Public Health Service contract HHSN-271-2013-00030, and the University of Maryland Brain Bank is a tissue repository of the National Institutes of Health (NIH) NeuroBioBank.

## Declaration of Competing Interest

The authors declare that the research was conducted in the absence of any commercial or financial relationships that could be construed as a potential conflict of interest.

## Notes

### Competing Interest Statement

The authors have declared no competing interest.

### Summary of Updates

During the assembly of this manuscript for upload, we inadvertently included incorrect figure legends for Figures 1-5; these have been corrected here. Additionally, new sequencing information resulted in the removal of one of the samples for Figure 1C and Table 1, which resulted in recalculation of the statistics and the number of samples ranked as outliers. Finally, the labeling/coloring of the groups in Supplemental Figure 1 in the last line of the legend was incorrect and this has been corrected. We apologize for these errors.

