## Supplemental Fig. 1. Pitolisant effects in male versus female MECP2Tg1 animals and wild-type (WT) littermate controls. for "Reciprocal regulation of the H_3_ histamine receptor in Rett syndrome and *MECP2* Duplication syndrome: implications for therapeutic development"

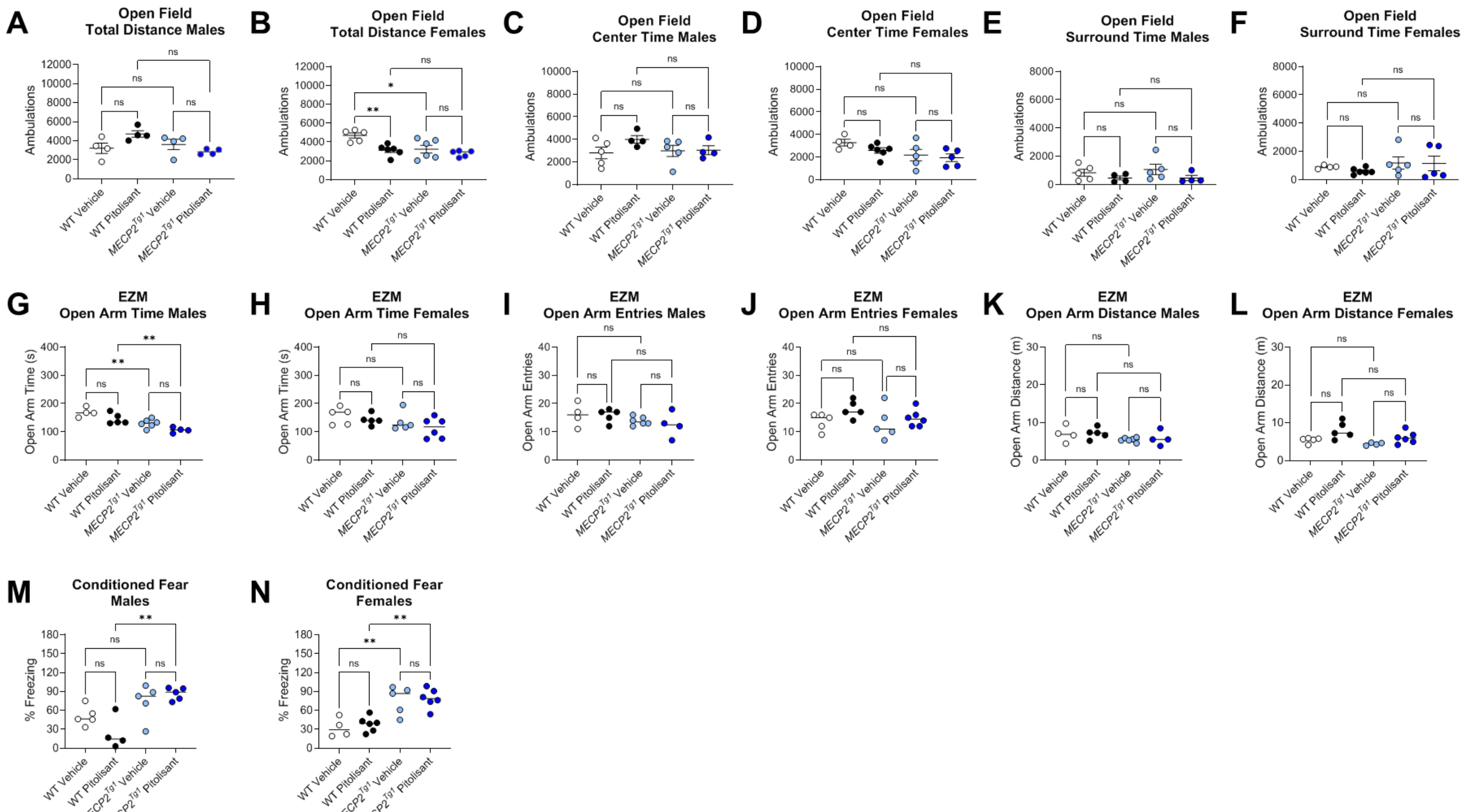


**Supplemental Fig. 1. Pitolisant effects in male versus female $MECP2^{Tg1}$ animals and wild-type (WT) littermate controls. First row: open field. (A)** Open field total distance traveled, males (1-way ANOVA, F (3, 12) = 3.459, p = 0.0512). **(B)** Open field total distance traveled, females (1-way ANOVA, F (3, 18) = 7.253, **p = 0.0022; Tukey's post hoc tests, WT vehicle (white) versus $MECP2^{Tg1}$ vehicle (light blue), *p = 0.0155; WT vehicle (white) versus $MECP2^{Tg1}$ pitolisant (dark blue), **p = 0.0023). All other comparisons were not significant. **(C)** Open field center time, males (1-way ANOVA, F (3, 14) = 1.279, p=0.3199). **(D)** Open field center time, females (1-way ANOVA, F (3, 16) = 2.385, p = 0.1074). **(E)** Open field surround time, males (1-way ANOVA, F (3, 14) = 1.228, p=0.3366). **(F)** Open field surround time, females (1-way ANOVA, F (3, 16) = 0.6851, p=0.5741). **Second row: Elevated zero maze. (G)** EZM open arm time, males (1-way ANOVA, F (3, 15) = 11.57, ***p = 0.0003; Tukey's post hoc tests, WT vehicle (white) versus $MECP2^{Tg1}$ vehicle (light blue), **p = 0.0076; WT vehicle (white) versus $MECP2^{Tg1}$ pitolisant (dark blue), ***p=0.0002; WT pitolisant (black) versus $MECP2^{Tg1}$ pitolisant (dark blue), **p=0.0096). All other comparisons were not significant. **(H)** EZM open arm time, females (1-way ANOVA, F (3, 17) = 1.885, p = 0.1705). **(I)** EZM open arm entries, males (1-way ANOVA, F (3, 15) = 1.231 p = 0.3331). **(J)** EZM open arm entries, females (1-way ANOVA, F (3, 17) = 1.663, p = 0.2126). **(K)** EZM open arm distance, males (1-way ANOVA, F (3, 15) = 1.470, p = 0.2627). **(L)** EZM open arm distance, females (1-way ANOVA, F (3, 16) = 4.804, *p = 0.0143; Tukey's post hoc test, WT pitolisant (black) versus $MECP2^{Tg1}$ vehicle (light blue), *p=0.0115). **Third row**: Conditioned Fear (CF). **(M)** CF, males (1-way ANOVA, F (3, 15) = 7.647, **p = 0.0025; Tukey's post hoc test, WT pitolisant (black) versus $MECP2^{Tg1}$ vehicle (light blue), *p=0.0133; WT pitolisant (black) versus $MECP2^{Tg1}$ pitolisant (dark blue), **p=0.0023). All other comparisons were not significant. **(N)** CF, females (1-way ANOVA, F (3, 17) = 11.35, ***p = 0.0003; Tukey's post hoc test, WT vehicle (white) versus $MECP2^{Tg1}$ vehicle (light blue), **p=0.0053; WT vehicle (white) versus $MECP2^{Tg1}$ pitolisant (dark blue), **p=0.0022; WT pitolisant (black) versus $MECP2^{Tg1}$ vehicle (light blue), **p=0.0067; WT pitolisant (black) versus $MECP2^{Tg1}$ pitolisant (dark blue), **p=0.0025). All other comparisons were not significant. Statistical comparisons represented on graphs are WT vehicle (white) versus $MECP2^{Tg1}$vehicle (light blue), WT pitolisant (black) versus $MECP2^{Tg1}$ pitolisant (dark blue), WT vehicle (white) versus $Mecp2^{+/+}$ pitolisant (black), and $MECP2^{Tg1}$ vehicle (light blue) versus $MECP2^{Tg1}$ pitolisant (dark blue).
